## Supplementary Materials for "The sperm hook in house mice: a functional adaptation for migration and self-organised behaviour"

**This file includes:**

Figures S1 to S3
Tables S1 to S3
Legends for Movies S1 to S10

**Other supporting materials for this manuscript include the following:**

Movies S1 to S10

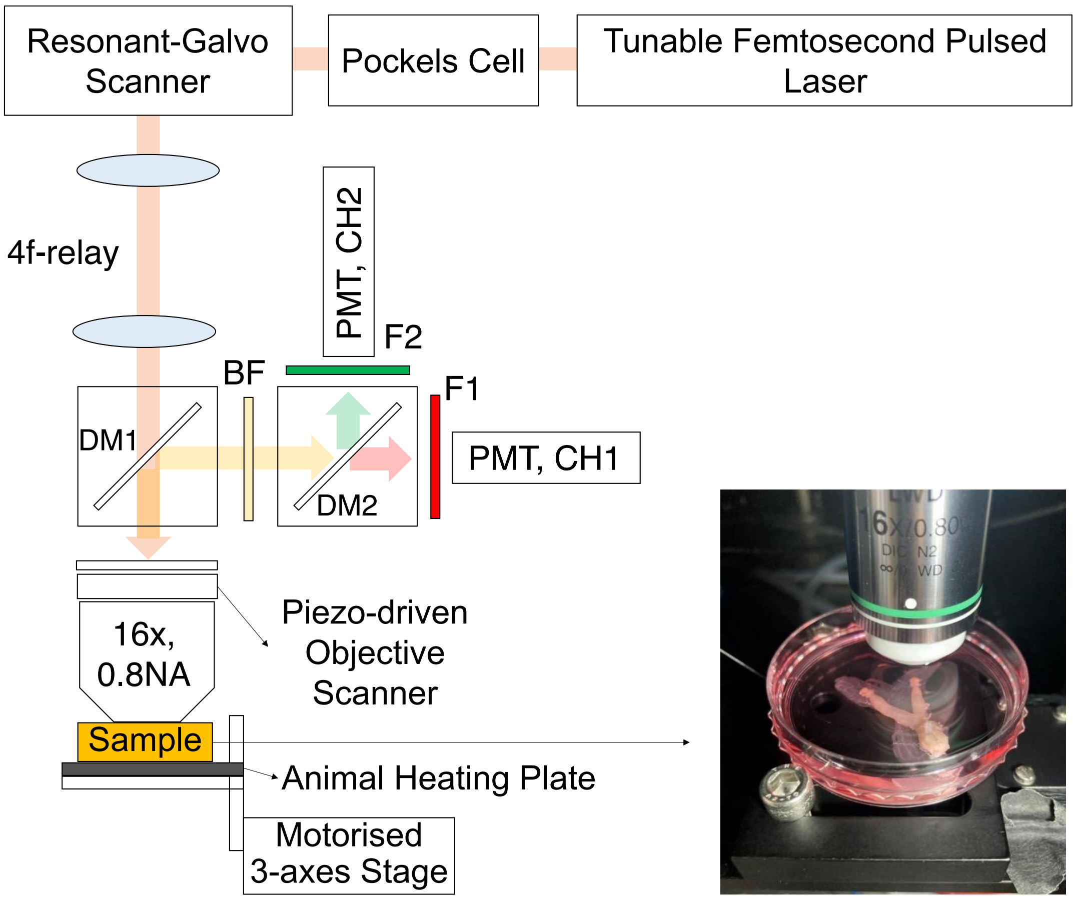


**Fig. S1.** Schematic diagram of the custom-built 2PSLM.

DsRed and eGFP were excited nonlinearly using a tunable high peak power femtosecond laser (Chameleon discovery/Coherent) and a water-immersion objective lens (CFI75 LWD 16X W/Nikon). The fluorescence emitted was collected by the same objective lens and detected by a pair of GaAsP photomultiplier tubes (PMT, H10770PA-40/Hamamatsu). Dichroic mirrors and filters; DM1(T735lpxrxt-UF3/Chroma), DM2(T565lxr/Chroma), BF(ET720SP-2P8/Chroma), F1(ET605/70m/Chroma), and F2(ET525/70m/Chroma) were used to split the excitation beam and emission light. A resonant-galvo scanner (RESCAN-GEN/Sutter instrument) enabled real-time fluorescence imaging of sperm behaviour at a speed of 30 frames per second for 512 pixels per line acquisition. A Pockels cell (M350-80-LA-02 KDP/Conoptics) allowed for rapid control of the laser beam intensity, homogenising the illumination across the field of view, and applying varying laser power per tissue depth. The sample position in all three dimensions was controlled by a Piezo-driven objective scanner (P-725.4CA/PI) and a motorised 3-axes stage (3DMS/Sutter instrument). A small animal heating plate (HP-4M/Physitemp) maintained the warmth of the mouse female reproductive tract at body temperatures.

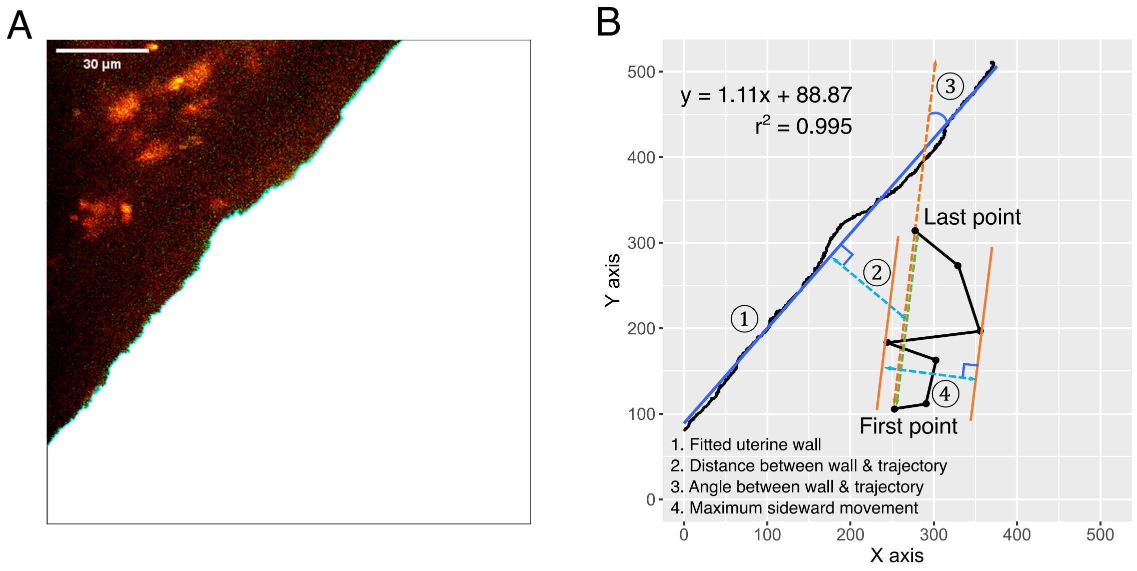


**Fig. S2.** Uterus wall and parameters that were used to measure sperm migration speed and linearity.

(**A**) Areas along the uterine wall that were relatively straight were selected. The boundary of these areas was identified using the object selection tool in Adobe Photoshop CC (23.1.0 version). (**B**) The uterine wall was approximated as a linear line using linear regression (①). The distance between a spermatozoon and the uterine wall was defined as the minimum distance between the midpoint of the track displacement and a sperm trajectory (②). The angle between sperm trajectories and the uterine wall was calculated as the angle (in radians) between the uterine wall (approximated line) and the straight line that connected the first and last points of a sperm trajectory – track displacement line (③). The maximum sideward movement was determined as the greatest distance between the parallel lines that aligned with the track displacement line at the positions of the sperm trajectory (④). SWR was then computed by dividing the track displacement by the maximum sideward movement.

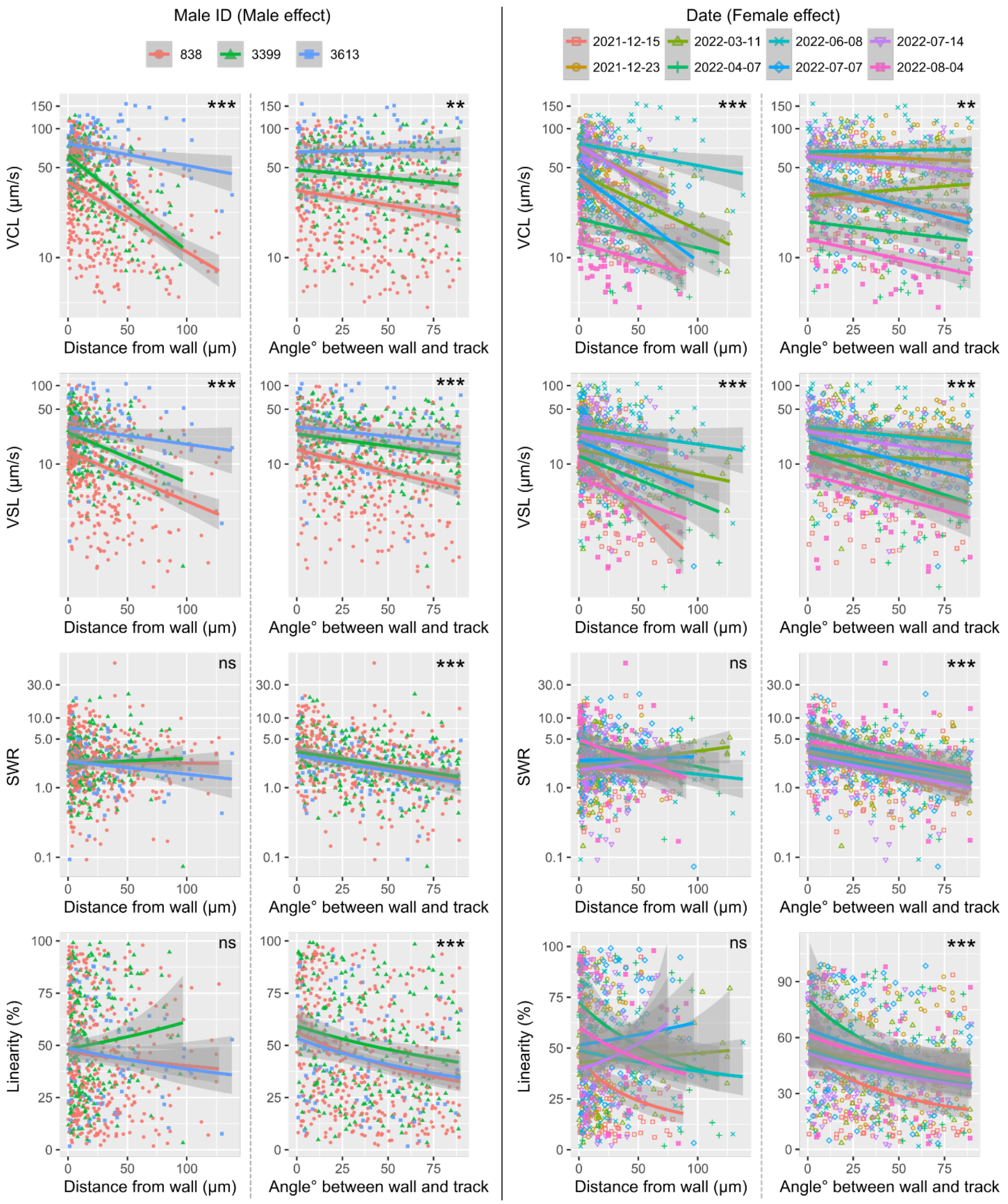


**Fig. S3.** Visualisation of male and female effects (two random effects) on sperm migration kinematic parameters. The model-fitted lines, separated by male ID and date (female), in the non-significant models, indicated as ‘ns’, show non-significant effects of the distance (SWR) or an inconsistent effect of the distance (Linearity) on the kinetic parameters.

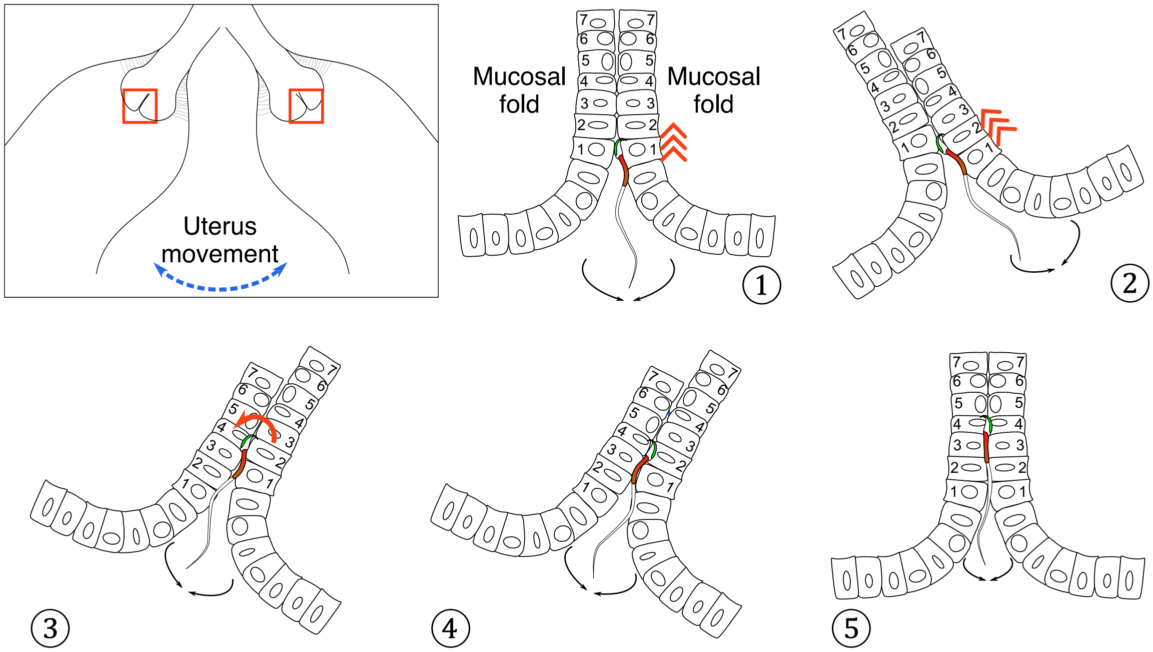


**Fig. S4.** A hypothetical model for sperm migration from the uterus to UTJ.

The upper left inset represents a uterine horn that moves to the left or right due to muscle contraction (exaggerated for visualization). The 5 subfigures with numbering represent zoom-in of the two red square frames in the inset. When the uterine horn moves from the centre to the right (① to ②), two facing surfaces between the two mucosal folds slide against each other. This sliding results in opening space where sperm can ascend – note that the sperm moves from cell 1 to cell 2 of the right mucosal fold (②). When the uterine horn moves from right to left (③), the two surfaces between the mucosal folds slide in opposite directions where the sperm can now reach cell 4 of the left mucosal fold. If sperm can turn over, its head can be attached to cell 4 of the left mucosal fold (④ to ⑤). Repetition of these procedures will make the sperm finally pass the CT and migration through narrow gaps between mucosal folds in intramural UTJ. This process appears to be as if sperm may slide through the space between mucosal folds when the space is too small for normal beating (Movie S5C).

**Table S1.** Basic information of 4 males that were used for the mating experiment.

| Male ID | Date of Birth (DOB) | Genotype | Note |
|---|---|---|---|
| A (No. 838) | 2021.04.21 | RBGS-CX3CR1 | |
| B (No. 3399) | 2021.08.17 | RBGS | |
| C (No. 3613) | 2021.09.01 | RBGS | |
| D (No. 3610) | 2021.09.01 | RBGS-CX3CR1 | Vasectomized male |

RBGS: CAG/Su9-DsRed2 and Acr3-EGFP, RBGS-CX3CR1: CAG/Su9-DsRed2, Acr3-EGFP, and Cx3cr1

**Table S2.** Mating records and the information of the females for sperm tracking.

| Date of mating | DOB (female) | Age (female) | Male ID |
|---|---|---|---|
| 2021-12-15 | 2021-09-01 | 15.0 weeks | A |
| 2021-12-23 | 2021-07-02 | 24.9 weeks | B |
| 2022-03-11 | 2021-05-10 | 43.6 weeks | A |
| 2022-04-07 | 2021-08-17 | 33.3 weeks | A |
| 2022-06-08 | 2021-10-17 | 33.4 weeks | C |
| 2022-07-07 | 2022-02-14 | 20.4 weeks | C |
| 2022-07-14 | 2021-10-17 | 38.6 weeks | A |
| 2022-08-04 | 2022-02-14 | 24.4 weeks | A |

Male ID: males (Supplementary Table 1) who mated with the subject females on the date.

**Table S3.** Summary results of the generalized linear mixed models (GLMM).

Each model represents sperm trajectory parameters that were log-transformed. In all models, we examined the effect of sperm to uterine wall distance (Distance from wall), angle between a sperm trajectory and uterine wall (respective angle with wall) and cropping of the acquired image (O: cropped vs X: uncropped). The GLMM for SWR showed a boundary (singular) fit warning message. However, two models that omitted one of random variables (Male or Date) did not result in any significant changes in the predictor variables ($p > 0.05$).

| Model | Predictors | Estimates | SE | *t* | *P* | 95% CI |
|---|---|---|---|---|---|---|
| VCL | (Intercept) | 4.0645 | 0.4189 | 9.7028 | 0.001** | 3.198, 4.921 |
| | Distance from wall | -0.0094 | 0.0007 | -12.8449 | <0.001*** | -0.011, -0.008 |
| | Angle with wall | -0.0019 | 0.0007 | -2.6255 | 0.009** | -0.003, 0 |
| | Cropped (O: X) | -0.2576 | 0.4368 | -0.5897 | 0.584 | -1.155, 0.652 |
| VSL | (Intercept) | 3.2745 | 0.4277 | 7.6558 | 0.002** | 2.434, 4.113 |
| | Distance from wall | -0.0104 | 0.0014 | -7.4718 | <0.001*** | -0.013, -0.008 |
| | Angle with wall | -0.0081 | 0.0014 | -5.9303 | <0.001*** | -0.011, -0.006 |
| | Cropped (O: X) | -0.0822 | 0.4202 | -0.1957 | 0.853 | -0.902, 0.716 |
| LIN | (Intercept) | -0.8236 | 0.1194 | -6.8999 | 0.001** | -1.058, -0.581 |
| | Distance from wall | -0.0009 | 0.0011 | -0.8632 | 0.388 | -0.003, 0.001 |
| | Angle with wall | -0.0061 | 0.0010 | -5.8469 | <0.001*** | -0.008, -0.004 |
| | Cropped (O: X) | 0.1701 | 0.1365 | 1.2459 | 0.273 | -0.113, 0.444 |
| SWR | (Intercept) | 1.1507 | 0.1458 | 7.8917 | <0.001*** | 0.732, 1.430 |
| | Distance from wall | -0.0003 | 0.0013 | -0.2149 | 0.830 | -0.003, 0.002 |
| | Angle with wall | -0.0108 | 0.0013 | -8.2828 | <0.001*** | -0.013, -0.008 |
| | Cropped (O: X) | 0.1566 | 0.1720 | 0.9107 | 0.399 | -0.384, 0.835 |

VCL: curvilinear velocity, VSL: straight-line velocity, LIN: linearity of forward progression, SWR: straight line-to-sideward movement ratio, ***: $p < 0.001$, **: $p < 0.01$, *: $p < 0.05$.

**Captions for Movie S1 to S10**

**Movie S1.**

Sperm migration in the uterus. Imaging was conducted around the area labelled as ‘Lumen’ in Figure 1A. (**A**) Most spermatozoa within the uterine volume are moving back and forth following the flow in the uterus. (**B**) Spermatozoa near the uterine wall (uterine epithelium) are more active and swim faster than those in the centre of the lumen. Sperm trajectories for the analysis in Figure 2 were also present later in the movie. Trajectories with different colours correspond to VCL of each trajectory (Blue: slow, Red: Fast). (C) Vertical scan imaging in different depths shows that most spermatozoa at the uterine wall (depth: 0 μm) are active and fast-moving. The further away from the uterine wall, e.g., depth: 30 μm, there are increasing number of inactive or slow-moving spermatozoa.

**Movie S2.**

The sperm hook helps sperm to determine migration directions in the uterus. Imaging was conducted around the area labelled as ‘Wall’ in Figure 1A. (**A**) When sperm reach the uterine wall, sperm change their migration direction. Most sperm change their heading direction in a way that their apical hook faces the uterine wall (pro-wall-hook direction) which allows sperm migrate straighter along the wall. In contrast, a few sperm exhibit an opposite heading direction such that their hook faces the uterine lumen (anti-wall-hook direction). This heading direction usually results in a departure of the sperm from the wall. (**B**) Sperm are tapping along the epithelium with their hook while migrating along the uterine wall.

**Movie S3.**

Sperm use their hook as an anchor to be attached to the uterine epithelium. Imaging was conducted around the area labelled as ‘CT (UTJ Entrance)’ in Figure 1A. (**A**) Sperm use their hook to be attached to the uterine epithelium (sperm hooking behaviour for anchoring). The apical sperm hook also helps sperm squeeze through other sperm in confined space. (**B**) Unattached (unanchored) or loosely attached sperm may be more easily squeezed out by uterine muscle contraction or fluid flow.

**Movie S4.**

The entrance of intra-mural UTJ (or CT) in the uterus has small spacing (almost closed inter-fold gaps) for mouse sperm to pass through. Imaging was conducted after excising and clearing the ‘Intramural UTJ’ part in Figure 1A. (**A**) These closed inter-fold gaps continue for about 100 μm from the entrance of UTJ. (**B**) The existence of a copulatory plug or mating history does not influence the opening of the inter-fold gap at the UTJ entrance (UTJ entrance seen from an orthogonal perspective with respect to **A**). (**C)** Even after mating with a male with working sperm, width of inter-fold gaps does not considerably change. Only a few sperm can pass through the inter-fold gap at a time due to its narrow width. Scanning direction of the intra-mural UTJ to get images is indicated with an arrow at the right upper corner in each Movie.

**Movie S5.**

Sperm behaviours and the movement of mucosal folds at the entrance of UTJ (CT). Imaging was conducted around the area labelled as ‘CT (UTJ Entrance)’ in Figure 1A. (**A**) There is no upsuck-like passive sperm transfer from the uterus to UTJ. Some unanchored sperm are sometimes pulled off by muscle contraction (peristaltic movement) in the intramural UTJ (indicated by an arrow).

(**B**) Two facing mucosal folds sometimes move in an opposite direction which causes the two mucosal folds to slide against each other. Sperm may use this moment to enter UTJ from the uterus. (**C**) As the sperm head is round enough despite its apical sperm hook, sperm can move forward (head direction) by sliding through a narrow lumen between mucosal folds. However, due to the sperm hook shape (anchor), it will not be easier for sperm to move backward (tail direction).

**Movie S6.**

Sperm unidirectional re-arrangement in a sperm cluster at a uterine crypt. Imaging was conducted around the area labelled as 'CT (UTJ Entrance)' in Figure 1A. (**A**) Asymmetrical sperm head shape may facilitate sperm unidirectional re-arrangement in sperm clusters at uterine crypts and CT. The unidirectional sperm clustering then results in synchronised sperm beating that pushes out other sperm by generating fluid flow or by beating other sperm directly. (**B**) Re-arranged sperm in a cluster sometimes move together to the same direction.

**Movie S7.**

An enormous unidirectional sperm cluster at CT in the uterus exhibits synchronised sperm beating. Imaging was conducted around the area labelled as 'CT (UTJ Entrance)' in Figure 1A. The synchronised sperm beating may prevent or hinder other sperm from approaching the UTJ entrance (CT) by pushing out approaching sperm.

**Movie S8.**

Accumulated sperm (sperm trains) are not found to swim faster than unlinked individual sperm in the uterus. Moreover, a large sperm accumulation cannot pass the narrow luminal space near the UTJ entrance (CT). These results suggest a disadvantage of sperm trains in sperm migration from the uterus to UTJ. Imaging was conducted around the area labelled as 'Wall' and 'Lumen' in Figure 1A.

**Movie S9.**

Various sperm behaviours in UTJ. Imaging was conducted around the area labelled as 'UTJ' in Figure 1A. (**A**) Sperm interact with the UTJ epithelium using their hook in various ways. They put their hook into a gap (crypt) and exhibit tapping- and stroking-like behaviour while they migrate through UTJ. (**B**) Sperm can more easily migrate when UTJ lumens get wider. Attached (anchored) sperm also beat faster when UTJ lumens get wider. (**C**) Dead or inactive sperm are accumulated in UTJ and may damage live sperm through collisions or hinder sperm migration in UTJ.

**Movie S10.**

The beating rate of the attached (anchored) sperm in UTJ changes over time. The speed of fluid flow and the luminal width of UTJ may be related to the beating rate. Imaging was conducted around the area labelled as 'UTJ' in Figure 1A.
